# *InVitro* Biocompatibility Evaluation of Acellular Porcine Dura mater Grafts and Native Dura

**DOI:** 10.1101/2023.04.24.538089

**Authors:** Ashma Sharma, Erika Moore, Lakiesha N. Williams

## Abstract

Damage to the dura mater may occur during intracranial or spinal surgeries, which can result in cerebrospinal fluid leakage as well as other potentially fatal physiological changes. As a result, biological scaffolds derived from xenogeneic materials are typically used to repair and regenerate dura mater post intracranial or spinal surgeries. The extracellular matrix of xenogeneic dura scaffolds has been shown to exhibit better cell infiltration and regeneration than synthetic material. In this study, we investigated the biocompatibility of native and decellularized porcine dura. Cell proliferation, cell viability, and mechanical properties of dural grafts were evaluated post re-seeding on days 3,7, and 14. Live-dead staining and resazurin salts were used to quantify cell viability and cell proliferation, respectively. Micro indentation was conducted to quantify the mechanical integrity of the native and acellular dura graft. The results show that the acellular porcine dural graft provides a favorable environment for rat fibroblast cell infiltration. Cell viability, proliferation, and micro indentation results on the acellular grafts are comparable with the native control porcine dura tissue. In conclusion, the porcine scaffold material showed increasing viable cells at each time point. The mechanics and biocompatibility results provide promising insight into the potential use of porcine dura in future cranial dura mater graft applications.

## Introduction

Cranial dura mater can be damaged during intracranial surgery or traumatic injury resulting in cerebrospinal fluid leakage, infection, or poor wound healing. These issues warrant dura replacement with either a biological or a synthetic graft. Despite extensive research, problems with dura replacement grafts include cerebrospinal fluid leakage, graft integration, and handleability [1][2]. Additionally, the suitability of the material being considered for the scaffold design is also governed by its biocompatibility, mechanical integrity, biodegradability, and scaffold architecture [1]. The most commonly used dura scaffold materials available on the market are absorbable synthetic material, allografts, and autologous fascia [2]–[7]. It has been shown from *in vitro* testing that after re-seeding, the cells can survive in the scaffold and release several bioactive molecules for regeneration [8]–[10].

Biological scaffolds derived from xenogeneic material are often used in various tissue engineering applications as their complex extracellular matrix (ECM) enhances constructive remodeling [11]. Some functions of the ECM include providing structural support for cells, contributing to enhanced mechanical properties, providing bioactive cues, acting as a reservoir of growth factors and their actions, and supporting tissue remodeling [12]. These xenogeneic decellularized ECM scaffolds are commonly isolated from small and large animals and have been used as a powerful tool in medical applications with minimal to non-immunogenic host responses [13][14]. Collagen, which contains non-allergenic, non-mutagenic, and non-migratory properties are the contributing factors for use of xenogeneic scaffolds as it is the main component of the ECM [15][16]. These collagen-based grafts are used as a reliable treatment for wound healing due to the amount of cell matrix interaction proteins [17][18].

Preclinical and clinical studies on bone tissue healing, epithelial skin and cornea regeneration support excellent biocompatibility with the use of collagen-based materials [13][17]–[22]. Xenografts, allografts, autografts, and synthetics are all options used to reconstruct both spinal and cranial dural defects [24]–[27]. Surgical reconstruction is determined by the amount of cerebrospinal fluid (CSF) leakage, location, size, and shape of the defect. Some of the problems with current grafts is the occurrence of CSF leaks, cost, availability, and impaired wound healing [26]. Each graft type has its own unique benefits. For example, autograft and allograft are less prone to infection, but the cost is high and availability is limited [27]–[29]. On the contrary, synthetic materials are easily available, but often initiate adverse immunogenic reactions, and offer inconsistent mechanics, which are not ideal [31][32]. Similarly, unforeseen immune reactions and infections have been observed after the use of biological scaffolds like cadaveric dura or xenograft pericardium [24][32]. Fibrous and fibromuscular soft tissue grafts are the most common material for dural covering, which should be viable and easily available. Research shows that decellularized ECM, in general, improves the growth and regeneration of tissues by allowing for fewer complications during repair [12]. Therefore, we propose exploring the use of decellularized porcine dura as a cranial dura replacement, which has not previously been considered.

In this study, we evaluate a decellularized porcine dura mater scaffold as a replacement for cranial dura through a series of *in vitro* tests as explained in experimental methods. Here we use a decellularized porcine dura model as a scaffold and seed it with primary rat fibroblast cells. The main objective of our work is to quantify cell viability, cell proliferation, and the mechanical integrity of decellularized porcine dura mater grafts compared to native porcine dura mater.

## Experimental Methods

### Decellularization and Cell Seeding

Dura mater was harvested from adult male pigs (Yorkshire, 6 to 8 months old) at a post-mortem time of 3 hours, at the large animal laboratory at the University of Florida Meat Processing Center (IACUC Protocol #201810442). Porcine dura mater was extracted within 2 hours of sacrifice and then washed with phosphate-buffered saline (PBS) solution to remove excess blood and debris. After the removal and cleaning process, the tissue was then cut into circular dura grafts of 14mm in diameter followed by decellularization processing using sodium dodecyl sulfate (SDS) solution. The tissues were decellularized for 48 hours in the solution and then washed with 1x PBS for 24 hours in a belly dancer. After washing, the tissue was constantly washed with ethanol and sterile saline alternating for three times. The decellularized dura was further sterilized with 2% peracetic acid (PAA) and ethanol: 2% peracetic acid/100% ethanol/dH2O (ratio v/v/v= 2/1/1) for 2-4 hours, followed by multiple washes by sterile PBS to remove residual acid. The decellularization technique was verified using a DNeasy (QIAGEN^®^, Germany) kit and a Nanodrop Spectrophotometer (Thermo Fisher Scientific, USA). The decellularized tissue and native tissue were freeze dried in liquid nitrogen and the dry weight of tissue samples was measured for DNA content normalization. The nanodrop quantification was done for n=6 animals for both decellularized and native tissues. The sterility was evaluated by incubating the sample in a cell culture medium (10% fetal bovine serum (FBS), 1% antibiotic antimycotic, and Dulbecco’s modified eagle medium (DMEM)) at 37°C. Also, some medium was dispensed in the empty wells as a control. Samples were assessed daily, and any color changes were taken as an indicator of contamination.

Once the sterility was confirmed, decellularized 14 mm diameter tissue grafts were placed in the same 24-well plate used for cell seeding. Tissue was emerged in the cell medium for 24 hours prior to cell seeding. After 24 hours, they were removed from the medium and kept in the biological safety cabinet for approximately 1 hour to dry the excess medium for better cell attachment. In the cell seeding process, the mixture containing 2 X10 ^6^ rat lung fibroblast cells and 100 μl cell culture medium was added in a dropwise manner to the decellularized porcine dura graft. Furthermore, 1 ml of additional cell culture medium was added to the seeded dura grafts after one hour of the initial seeding process to ensure the possibility of cell attachment. In the current study, all testing and analysis were conducted at three specific time points: 3 days, 7 days, and 14 days after initial seeding, and the cell culture media was changed daily.

### Histological Analysis

Freshly isolated native, decellularized, and re-seeded dura mater of randomly assigned male pigs were fixed using formalin for all time points, embedded in paraffin, and sectioned. After that, the sectioned tissue was stained with hematoxylin & eosin (H&E) to confirm collagen fiber integrity, cell number density, and area fraction of collagen fibers. The microscopic images of the dura mater (n=3 per sample) were analyzed with ImageJ software (National Institutes of Health) to quantify the microstructure of the extracellular matrix (ECM) of the tissue. The structural integrity of collagen and elastin was also examined qualitatively to assess the microstructural alterations [33].

### Cell Viability

Cell viability at three-time points was determined by calceinAM and ethidium-homodimer. Cell-seeded constructs were incubated in serum-free media supplemented with 4mmol/l calceinAM and 2mmol/l ethidium-homodimer for one hour at 37°C. Constructs were then washed with PBS and covered with glass coverslips. After that, the construct was immediately scanned using a fluorescence microscope (Leica DFC 300 FX, Leica Microsystems, Wetzlar, Germany). The live cells were qualitatively analyzed as the number of green stained cells.

### Cell Proliferation

The cell proliferation assay was performed at three-time points (3, 7, 14 days) using resazurin stock solution with a concentration of 5 mM and was prepared by dissolving resazurin powder (resazurin sodium salt *CAS 62758-13-8*) into distilled water. Furthermore, the resazurin working solution was prepared by mixing 10% of the resazurin stock solution with the cell culture medium. The re-seeded dura graft samples for the three-time points were then incubated in a resazurin working solution for 4 hours at 37 °C with 5% CO_2_ atmosphere to convert resazurin to resorufin. Reduction percentage refers to the degree of reduction of the Alamar blue by proliferating cells. The higher percentage of reduction means the greater proliferation of cells in culture.

Triplicate aliquots of 100 mL of the incubated resazurin working solution were taken to measure fluorescent intensity using a microplate reader (SpectraMax M5/M5e) at a wavelength of 590 nm. Metabolic activity of the cells, given by the percent reduction of assay (PRA) was measured by using Eqn. (i), where plain resazurin working solution was taken as untreated control, the autoclaved solution was taken as 100% reduced, and absorbance values were calculated [25][26]

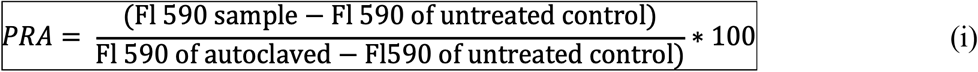

### Micro indentation

The samples were indented using a custom built piezo-electric stage (P-628.1 CD, Physik Instrumente) along with piezo amplifier (E-625.CR Servo Controller, Physik Instrumente) [36]. A custom-made spherical indenter (dia. of 8 micrometers) was used to indent the sample at five locations. A total of four samples were indented at each time points for the native (n=4 male pigs) and decellularized tissue (n=4 male pigs). All tests were performed at room temperature and PBS was sprayed for hydration. The total indentation depth for loading and unloading was 100 μm at a rate of 10 μm/s and a holding time of 60 s between loading and unloading [27][28].The Hertz contact model was used to determine the elastic modulus (E_Hertz_)using force as a function of indentation depth [38][37].

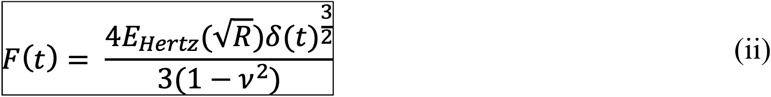

where F = force calibrated by stiffness and displacement, R is the radius of the tip, δ is the indentation depth, and ν is the Poisson’s ratio.

The Hertz contact model was then rearranged to determine the effective modulus as a function of time for soft samples in quasi-static conditions.

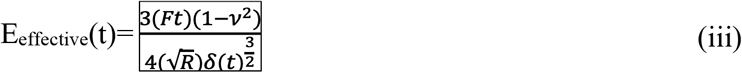

where R = 4 mm, δ is 100 μm and ν is 0.5.

The effective modulus at the equilibrium steady state modulus (SSM) is similar to the stress relaxation modulus when t = ∞. [39]

The equation (iii) was fitted to the relaxation phase of our indentation results using the MATLAB curve fit tool to calculate the steady-state modulus (E_p_) [36][38][40]. Further, the Goodness of Fit function in MATLAB was used to evaluate fits using NMSE whose values were above 0.8.

### Statistical Analysis

All experimental results were expressed in the form mean ± standard deviation. A one-way ANOVA followed by post-hoc test for multiple pairwise comparison was performed to evaluate statistical differences between the native, decellularized, and re-seeded dura mater at three time points. A value of p< 0.05 is assumed to be statistically significant in this study.

## Results

### Decellularization

A small thickness reduction was observed in the decellularized dura. The average thickness of native dura mater is 0.69 ±0.20 mm and 0.65 ±0.15 mm for decellularized dura mater. The thicknesses were not significantly different. The DNA content was analyzed using both a nanodrop spectrophotometer (Thermo Fisher Scientific, USA) along with a DNeasy kit (QIAGEN^®^, Germany). The DNA content of decellularized dura mater is less than 50ng per mg dry weight. The DNA content as shown via spectrophotometry is 423.33±69.58 ng/mg and 32.75 ±8.84 ng/mg for native and decellularized samples, respectively. An unpaired T-test for DNA content reflected significant differences in native and decellularized samples with a p-value of 0.009 (p<0.05).

### Histology

The histological images show the lateral view of dura mater after 3, 7, and 14 days of re-seeding as well as in native and decellularized conditions (**Fig. 1**). The H&E stained images show fiber morphology (pink area) of the dura mater. However, as expected, the fibroblasts (blue dots indicated by red arrows) were absent in the decellularized tissues, while they were observed in both the seeded and native dura. This indicates that the procedure used for decellularizing and re-seeding in the current study was appropriate.

**Figure 1:**
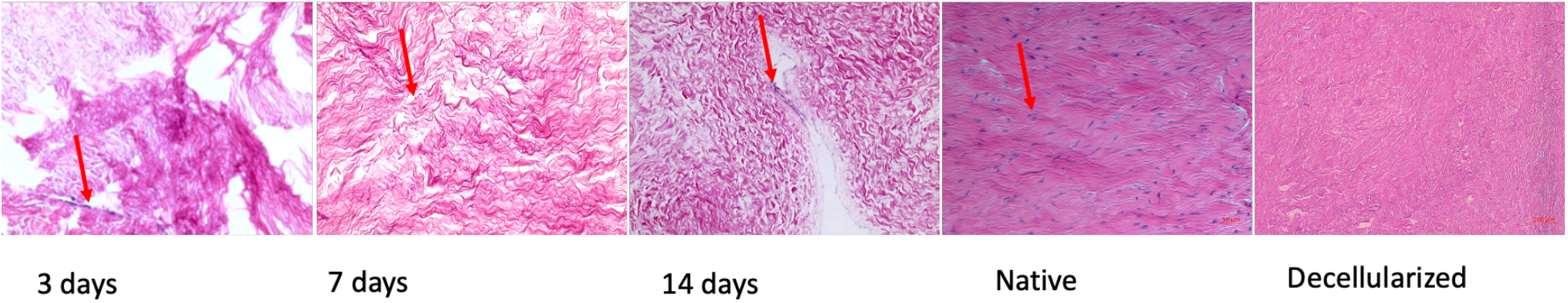
Histological images of dura mater in transverse direction at 20X magnification (scale Bar 100µm) at 3, 7, and 14 days after re-seeding compared with decellularized and native tissue (red arrow represents the presence of cells).

The area fraction of dura mater, which is taken as the ratio of the area of fibers to the total area was calculated to determine the effect of decellularization and re-seeding on the structure of dura mater. The area fraction of collagen represented by the histogram, as shown in **Fig. 2**, was calculated to be 0.897±0.046 for native tissue, 0.902±0.059 for decellularized tissue, 0.864±0.030 at 3 days, 0.902±0.028 at 7 days, and 0.916±0.001 at 14 days post re-seeding. The P-value determined from the one-way ANOVA conducted on the obtained data in each sample condition was greater than 0.05, indicating statistically insignificant results.

**Figure 2:**
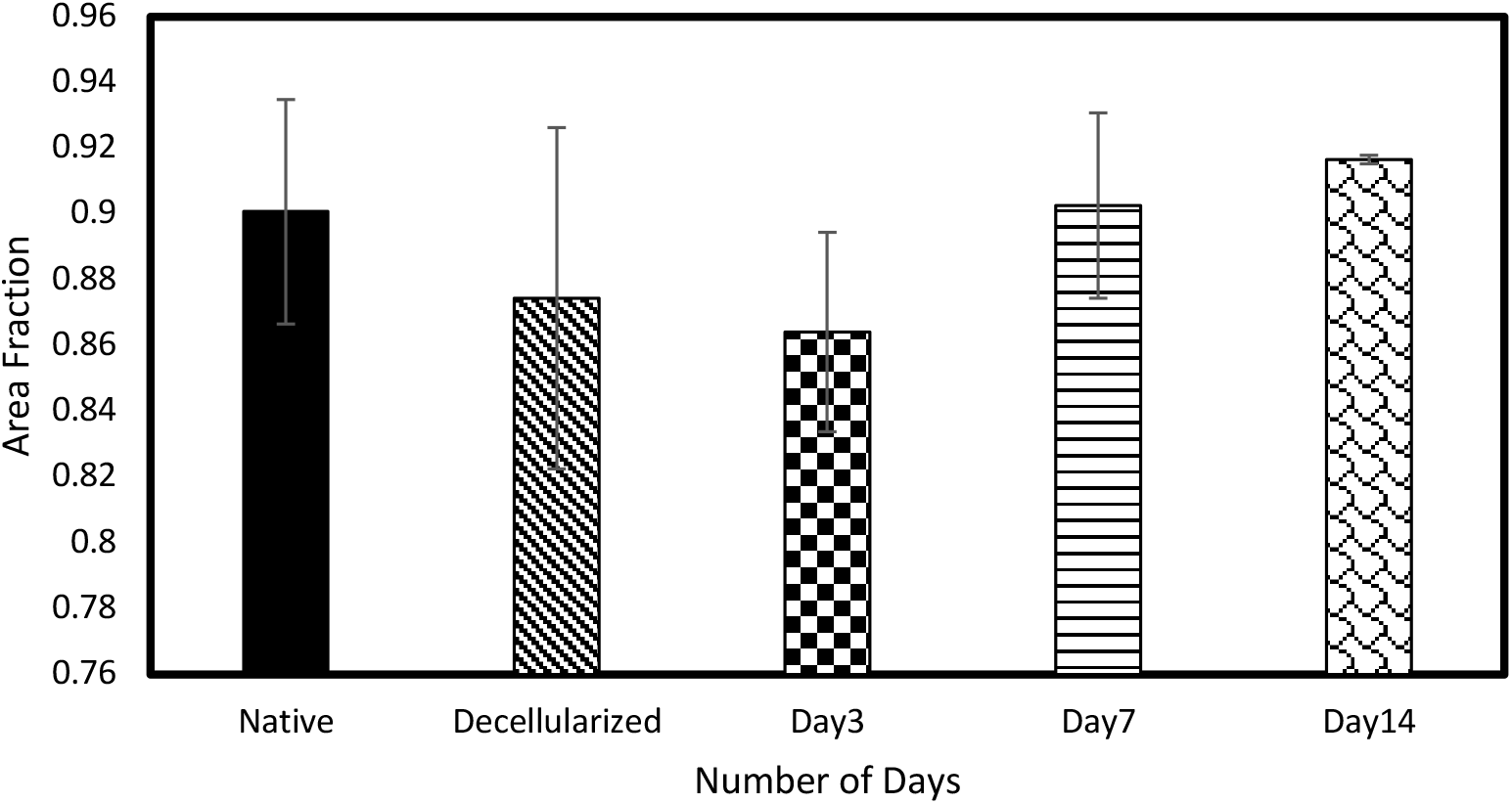
Bar graph of the area fraction of collagen fibers of native dura, decellularized dura, and re-seeded tissue at prescribed time points (3 days, 7 days and 14days) (no significant differences between groups).

**Fig 3** shows a consistent increase in cell count for native and re-seeded dura mater at three different time points. Based on the Image J analysis, the number of cells (H and E stained nuclei) were calculated as 90±6, 182±22, and 280±18 for the dura mater after 3, 7, and 14 days of re-seeding, respectively, and 492±9 cells in native condition. As intended, the cell count was observed to increase with the increasing number of days after the re-seeding process, while the native dura mater still had the highest number of cells. Furthermore, from the one-way ANOVA along with the post hoc test, the p-value was determined to be less than 0.05 for all different conditions, indicating that the cell count data obtained for dura mater at different days after re-seeding were statistically different (Table 1).

**Table 1:** P-value determined by post hoc test for all different conditions of cell count data obtained for dura mater at different days after re-seeding.

|  | Native | Day 3 | Day 7 | Day 14 |
| --- | --- | --- | --- | --- |
| Native | - | 0. | 0 | 0 |
| Day 3 | 0 | - | 0.0000229 | 0 |
| Day 7 | 0 | 0.0000229 | - | 0.0000123 |
| Day 14 | 0 | 0 | 0.0000123 | - |

**Table 2:** P values determined by post hoc test for proliferation using resazurin assay for dura mater at different days after re-seeding.

|  | 3 days | 7 days | 14 days |
| --- | --- | --- | --- |
| 3 days | - | 0.625 | 0.00142 |
| 7 days | 0.625 | - | 0.0018 |
| 14 days | 0.00142 | 0.0023 | - |

**Figure 3:**
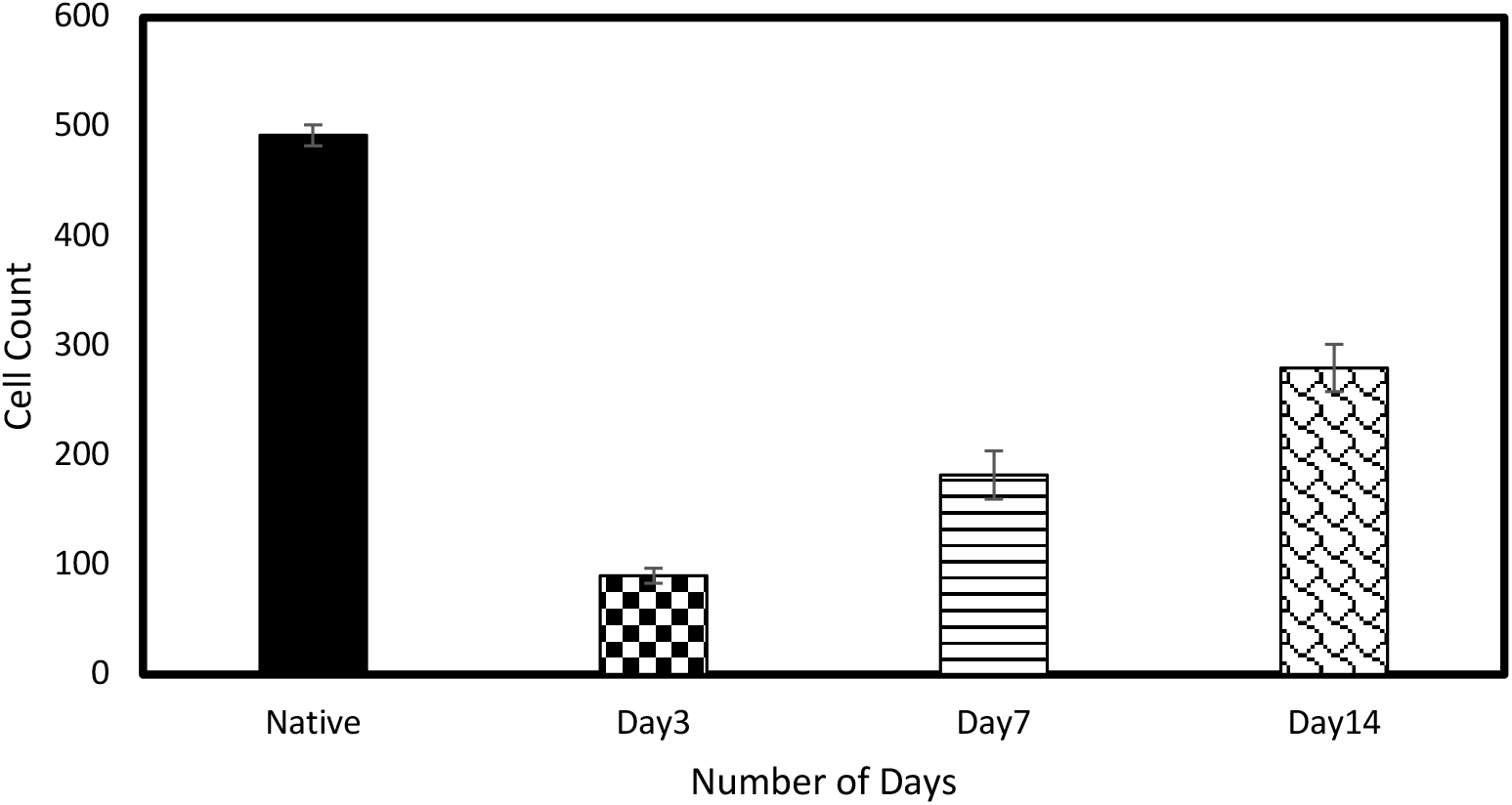
Bar graph of cell count of re-seeded dura mater at 3 days, 7 days and 14 days post re-seeding compared with native tissue (Location for cell count: four corners and two central areas).

### Cell Viability

Cell viability tests using live dead staining was performed to analyze the health and growth of cells in dura mater tissues after 3, 7, and 14 days of the re-seeding process. These results were compared with the one observed for native dura mater in **Fig. 4**. Green illuminated features shown in **Fig. 4** represents the live cells, while the red features represent dead cells. In all the images shown in **Fig. 4**, the presence of red (dead) cells was not evident and the cell number and cell proliferation was observed to increase as the time point increased in the case of re-seeded samples.

**Figure 4:**
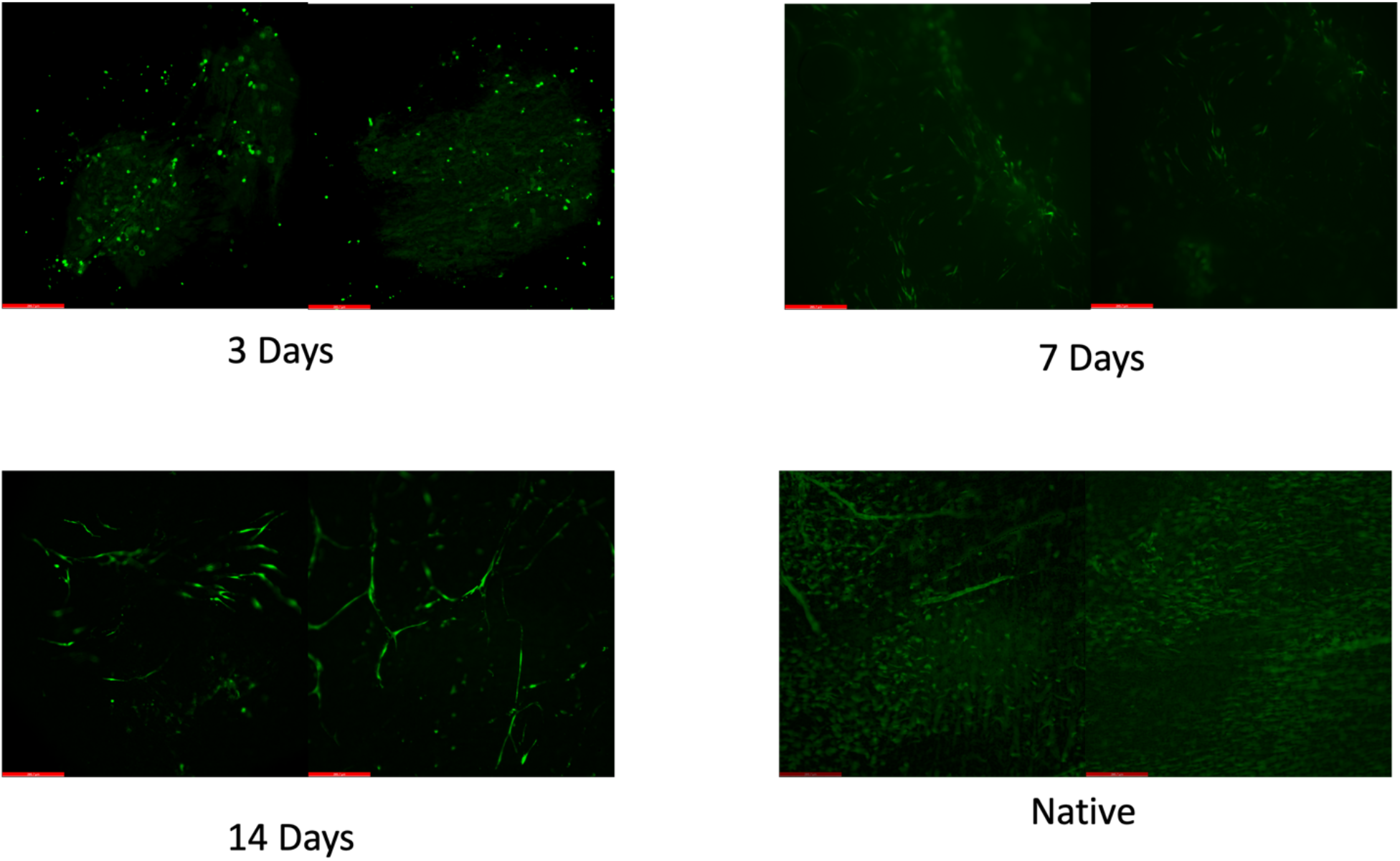
Live dead staining image of the fibroblast cells grown on the prepared sample after 3 days,7 days, and 14 days of incubation compared with native dura matter. Live cells are shown in green (scale Bar 266.7µm).

### Cell proliferation

To determine the metabolic activity of fibroblast cells, Percentage Reduction Assay (PRA) values were calculated using Eqn. (i) for re-seeded dura mater at the three different time points (3, 7, and 14 days). **Fig 5** shows the PRA values to be lower on days 3 and 7 compared to the value determined at day 14, which may be associated with higher metabolic activity. The amount of percentage reduction in the test well increased as the time points increased indicating presence of cell metabolic activity. Similar to all other analyses, one way ANOVA along with post-hoc test was also performed for the PRA values at different time points. The PRA values obtained for 3 and 7 days were not statistically different (P value of 0.3). On the other hand, results obtained for 3 and 14 days (P value of 0.0002) and 7 and 14 (P value of 0.002) after re-seeding were statistically different.

**Figure 5:**
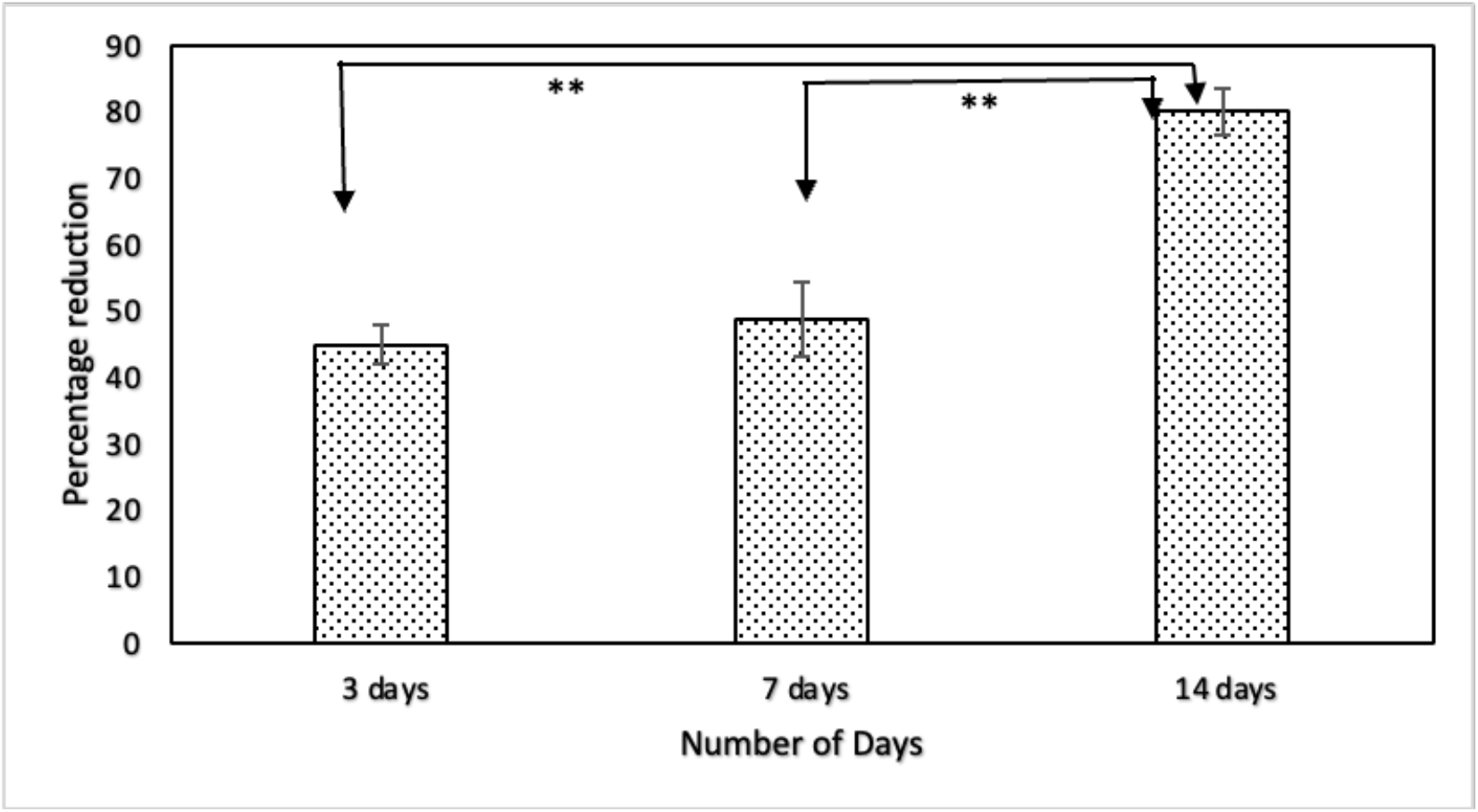
Proliferation of fibroblast cells on the prepared sample by using percentage reduction of resazurin assay using fluorescence.

**Figure 6:**
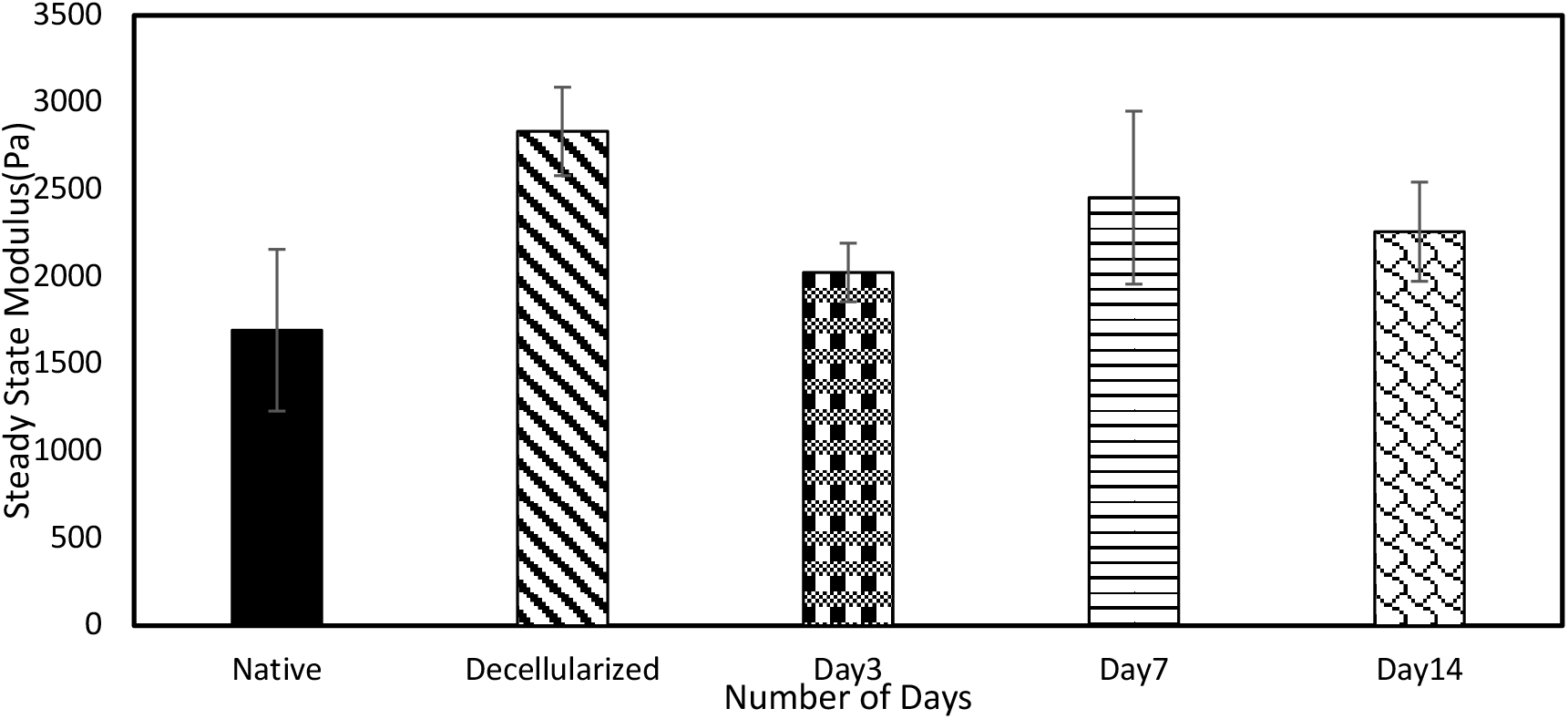
Bar Graph of steady state modulus from micro indentation at 3, 7, and 14 days after re-seeding on native and decellularized dura.

### Micro indentation

The modulus was calculated for native, decellularized and re-seeded samples for 3,7, and 14 days using the steady state moduli. The indentation force increased gradually during loading, decreased in response to holding, and decreased gradually during unloading. Towards the end of unloading, the force became negative in response to tissue adhesion and returned to its initial value upon indenter-sample separation [37]. Results show that the steady state modulus is slightly increased in decellularized dura as compared to the native dura. The average steady state moduli of native dura is 2127.75±15.20 Pa, 2725±237 Pa for the decellularized dura, 2025.7±168.18 Pa for the tissue 3 days after re-seeding, 2455.15±495 Pa for tissue 7 days after re-seeding, and 2259.46±284.51 Pa for tissue 14 days after re-seeding. The comparison of native, decellularized tissue and tissue post re-seeding using one-way ANOVA followed by post-hoc test did not show significant differences in micro indentation (p-value greater than 0.2).

## Discussion

The number of cells within the re-seeded dura graft proliferated with the increase in the number of days. The highest number of cells was seen in the sample after 14 days of re-seeding, while the lowest cell count was seen in the sample after 3 days of the re-seeding process (**Figs. 1 and 3**). In contrast, the change in the area fraction of collagen (**Fig. 2**) was observed to be statistically insignificant within decellularized, native, and re-seeded tissue, which may imply that the re-seeding process did not change the collagen amount in the microstructure of dura mater. Studies have shown that the chemical decellularization methods usually lead to certain destruction in fibers [41] and the chemical detergent with sodium dodecyl sulfate may compromise the fiber substrate [42][43]. However, the DNA content within the dura tissue after decellularization decreased by more than 90% and is less than 50ng per mg dry weight, which is adequate to satisfy the intent of decellularization [44]. Additionally, cell proliferation may also indicate that the decellularization process provides a suitable ECM, which facilitates cell attachment and promote tissue regeneration [45].

Fibroblasts in the ECM are very important to interpret the cellular activity, remodeling, and proliferation tendency. It is important to note that cell count analysis provides an estimation of the number of cells present within the sample, but do not indicate the viability. Hence, to determine the state (live or dead) of cells, a cell viability test was conducted using the live dead staining assay. The results presented in **Fig. 4** showed that a small number of scattered live cells were present at the superficial layer of the tissue after three days of the re-seeding process, while after seven days, the number of live cells increased along with some elongation. After 14 days of re-seeding, the cells were observed to be more elongated in shape and highly interconnected with the surface, akin to that of a fibroblast cell. The random and interconnected pattern of cells also indicated that the cells were infiltrated in the extracellular matrix up to 14 days post re-seeding. A similar pattern where cells were stable with good graft integration was also reported in a study by Elson et al., thus confirming graft suitability [46].

Metabolic activities such as cell viability, cytocompatibility of biomaterials, and cell proliferation were quantitively evaluated. Cytocompatibility tests using the resazurin-based assay was used because of their high sensitivity and cost-effectiveness. Resazurin, a cell-permeable and virtually nonfluorescent dye, is reduced to resorufin as a result of cellular metabolism (i.e., DNA content and glucose consumption) which changes its color in the process. Based on the change in color, the PRA was determined using Eqn. (i), which provided a quantitative measure of proliferation [35]. The percentage of reduction increased from day 3 to day 14, which is consistent with the number of cells being greater at 14 days after re-seeding as compared to 3 days and 7 days. However, the cell growth rate slowed after 7 days. Slow growth may be a result of the coverage of free area available on the scaffold and the lack of new surface area to grow as also indicated by Dan et al. [47]. The study by Ng et al. revealed a lack of linear correlation of reduction assay where there is the presence of high cell density, which supports the slower growth rate in this study after 7 days [48]. In this study, after 7 days, the cells could have gone into arrest after reaching a certain degree of confluency which might have caused apoptosis and thus a slow increase in the percentage of reduction [49].

Micro-indentation testing showed that the steady-state compressive modulus of the decellularized dura was similar to the native dura (native dura: 2127 ±15 Pa, decellularized dura:2725±237 Pa). Also, the one-way ANOVA reveals that the steady state modulus for three days, seven days and fourteen days post re-seeding were similar with P value greater than 0.05(3 days: 2025±168 Pa, 7 days: 2455±495 Pa, and 14 days: 2259±284 Pa). The compressive modulus calculated from micro indentation revealed that the mechanics of the native tissue was preserved after decellularization. Additionally, the mechanics were similar post re-seeding. The compressive modulus calculated from micro indentation is difficult to compare across soft tissue because the response depends on the indentation depth and loading rate [36]. We specifically controlled those in our experimental protocol. In soft matter, when the indentation rate increases it takes less time to dissipate energy, which eventually increases the stiffness [38]. When the thickness is inconsistent, the indentation rate changes the moduli. Loading in the compressive stress state locally under micro indentation is likely the reason for the resulting different moduli values from uniaxial global compression tests. Although the location was controlled, thickness variations and the number of cells across the dura mater may have caused some of the scatterings of the steady-state compressive modulus.

An effective cranial dural substitute should have various long-term and short-term characteristics. These include appropriate strength, a thickness similar to that of native tissue, minimal inflammation, minimal scarring, and minimal risk of transmission of infectious diseases [5][50]. The *in vitro* evaluation of the scaffold is a standard method to allow for direct comparison across the material. The regenerative process is very complex, and in this *in vitro* study, we do not attempt to mimic the condition for clinical applications. Nevertheless, our study successfully captures the features for tissue regeneration to lend insight into *in vivo* applications using decellularized porcine dura mater.

## Conclusions

In conclusion, this study showed that a xenogeneic scaffold derived from decellularized porcine dura has a similar architecture as native porcine dura. The microstructural analysis of collagen fibers confirms that the structural integrity of the graft remains intact over the prescribed time points. Here, a stable scaffold supported cell attachment and proliferation over 14 days. Future work will include *in vivo* subcutaneous testing using an animal model. This data will allow for assessing the feasibility of using the current porcine dura model as a future replacement for native dura in clinic.

